# GNG2–MRAS interaction regulates SHOC2-dependent RAF activation and MAPK signaling

**DOI:** 10.64898/2026.10.31.763144

**Authors:** Govinda R. Navale, Bamidele Towolawi, Colin Christoff, Sravani Malasani, Zhiwei Liu, Zhihong Wang

## Abstract

Dysregulation of the mitogen-activated protein kinase (MAPK) signaling pathway is a hallmark of many human cancers and RASopathies. The SHOC2–MRAS–PP1Cα (SMP) holophosphatase complex is a crucial regulatory node of this pathway. It activates the MAPK pathway by dephosphorylating the inhibitory phosphoserine on RAF kinases and thereby relieving RAF autoinhibition. Although GNG2, the γ-subunit of heterotrimeric G proteins, has recently emerged as a negative modulator of MAPK signaling, its molecular mechanism of action remains incompletely understood. Here, we identify GNG2 and its truncated variants as inhibitors of SMP complex assembly by directly targeting MRAS. Using NanoBiT live-cell bioluminescence, co-immunoprecipitation (Co-IP), biolayer interferometry (BLI), and molecular dynamics simulation studies, we demonstrate that full-length GNG2 (residues 1–71aa) and two helical fragments (Δ1, residues 1-29aa and Δ2, residues 30-71aa) bind MRAS, disrupt SMP complex formation, and therefore prevent RAF dephosphorylation. Expression of the truncated GNG2 variants significantly reduced phosphorylation of ERK in cells, consistent with inhibition of pathway activation. Furthermore, a short peptide synthesized from the GNG2 Δ2 sequence effectively disrupted SMP complex formation in both purified protein systems and live cells and potently attenuated MAPK signaling. Molecular dynamics simulation studies revealed that GNG2 variants engage with the Switch I/II regions of MRAS, sterically interfering with SHOC2–PP1Cα association. Together, these findings establish GNG2 as a direct regulator of SMP complex assembly and demonstrate that GNG2-derived peptides represent a promising strategy for disrupting SHOC2–MRAS interactions and suppressing MAPK signaling in RAS/RAF-driven pathologies.

## Introduction

The mitogen-activated protein kinase/extracellular signal-regulated kinase (MAPK/ERK) pathway is a highly conserved signaling cascade that regulates fundamental cellular processes, including proliferation, differentiation, survival, and migration (1, 2). Dysregulation of this pathway is a hallmark of many human diseases, such as cancers and RASopathies (3, 4). Central to this pathway are the RAF serine/threonine kinases (ARAF, BRAF, and CRAF), which function immediately downstream of RAS GTPases to relay signals from activated RAS to the MEK–ERK kinase cascade. Persistent activation of the RAS–RAF signaling axis promotes uncontrolled cell growth, malignant progression, and therapeutic resistance (5, 6). At the molecular level, RAF activation is governed by dynamic protein–protein interactions (PPIs), phosphorylation–dependent autoinhibition, membrane recruitment, and dimerization (7, 8).

A key regulatory checkpoint controlling RAF activation is the SHOC2–MRAS–PP1C (SMP) holophosphatase complex. The SMP complex activates RAF by catalyzing the dephosphorylation of conserved inhibitory phosphoserine residues (ARAF S214, BRAF S365, and CRAF S259) (9–12). In resting cells, phosphorylation of these sites promotes binding of 14-3-3 proteins, stabilizing RAF in an autoinhibited monomeric conformation that prevents formation of the kinase-domain dimer required for activation (13). Upon growth factor stimulation, GTP-bound MRAS recruits SHOC2 and PP1Cα to assemble the SMP holophosphatase complex at the plasma membrane. SHOC2 functions as a leucine-rich repeat (LRR) scaffold that simultaneously engages MRAS and PP1Cα, positioning the phosphatase to selectively dephosphorylate RAF. Removal of the inhibitory phosphoserine destabilizes the autoinhibited RAF–14-3-3 complex, promotes RAF membrane enrichment and dimerization, and enables downstream activation of the MEK–ERK signaling cascade (14). Recent structural and functional studies have established that SMP assembly is highly cooperative and depends on the active GTP-bound conformation of MRAS, making the complex a finely tuned regulatory switch for MAPK signaling (15–18).

Growing evidence further highlights the importance of SHOC2 in oncogenic signaling. Data from the Broad Institute’s Cancer Dependency Map and multiple independent studies demonstrate that tumors harboring oncogenic RAS mutations, particularly G12, G13, and Q61 variants, exhibit a strong dependence on SHOC2 for proliferation and survival (18, 19). Consistent with these observations, genetic ablation of SHOC2 selectively suppresses the malignant properties of RAS– mutant tumor cells and sensitizes them to MEK inhibition. Collectively, these findings identify the SMP complex as a critical signaling hub and an attractive therapeutic target for RAS-driven malignancies (15–18). Despite the emerging importance of the SMP complex in RAF activation, relatively little is known about endogenous proteins that directly regulate its assembly.

Because the SMP complex functions upstream of RAF activation, disrupting its assembly is a promising strategy to suppress MAPK signaling and potentially overcome resistance mechanisms associated with BRAF or MEK inhibitors (20–22). Unlike kinase inhibitors that target catalytic activity, modulation of protein–protein interactions within the SMP complex offers the opportunity to selectively interfere with the molecular events required for RAF activation. Consequently, chemical or peptide-based modulators of SMP complex assembly could serve as valuable tools for dissecting MAPK pathway regulation and as leads for therapeutic development.

GNG2, a γ subunit of heterotrimeric G proteins, forms a Gβγ–dimer together with a Gβ subunit (23) and participates in diverse signaling pathways regulating cell proliferation, differentiation, and angiogenesis (24). Increasing evidence supports a tumor-suppressive role for GNG2 in triple-negative breast cancer (TNBC) (25), colon cancer (26), and melanoma (27), where it inhibits proliferation and metastasis by attenuating MAPK (25), PI3K/AKT (28), and FAK (29) signaling. Interestingly, GNG2 has been reported to interact with the small GTPase MRAS, raising the possibility that it directly regulates SMP complex assembly and RAF activation (24, 25). However, the molecular basis of GNG2-mediated modulation of MAPK signaling remains largely unknown.

In the present study, we investigated the molecular mechanism by which GNG2 regulates MAPK signaling through the SMP complex. Guided by structural analyses, we generated two truncated helical variants of GNG2 and evaluated their ability to modulate SMP complex assembly and downstream signaling. Using live-cell NanoBiT complementation assays, co-immunoprecipitation, biolayer interferometry, molecular dynamics (MD) simulations, and biochemical analyses, we demonstrate that GNG2 directly binds the Switch I/II interface of MRAS, competitively disrupting SHOC2 binding and preventing assembly of the SMP holophosphatase complex. Consequently, GNG2 inhibits RAF dephosphorylation, stabilizes RAF autoinhibition, and suppresses ERK activation. Remarkably, a synthetic peptide derived from the C-terminal helical region of GNG2 retained potent inhibitory activity in both purified protein systems and live cells. Together, our findings identify GNG2 as a previously unrecognized regulator of SMP complex assembly and establish GNG2-derived peptides as promising molecular scaffolds for targeting SHOC2-dependent MAPK signaling in RAS-driven pathologies.

## Results

### GNG2 Suppresses MAPK Signaling through Interaction with MRAS

GNG2 is a 71-residue γ-subunit of heterotrimeric G proteins (28). AlphaFold2 predicts that GNG2 adopts two α-helices connected by a flexible, partially disordered region, suggesting that individual structural elements may contribute independently to protein interactions (**Figure 1A**). Guided by this structural model, we generated two truncated variants, GNG2 Δ1 (residues 1–29aa) and GNG2 Δ2 (residues 30–71aa) (**Figure 1B**). Co-immunoprecipitation confirmed that MRAS binds to GNG2 when Myc-MRAS and Flag-GNG2 FL were co-expressed in HEK293 cells (**Figure 1C & Figure S1**), consistent with a previous report (25).

**Figure 1.**
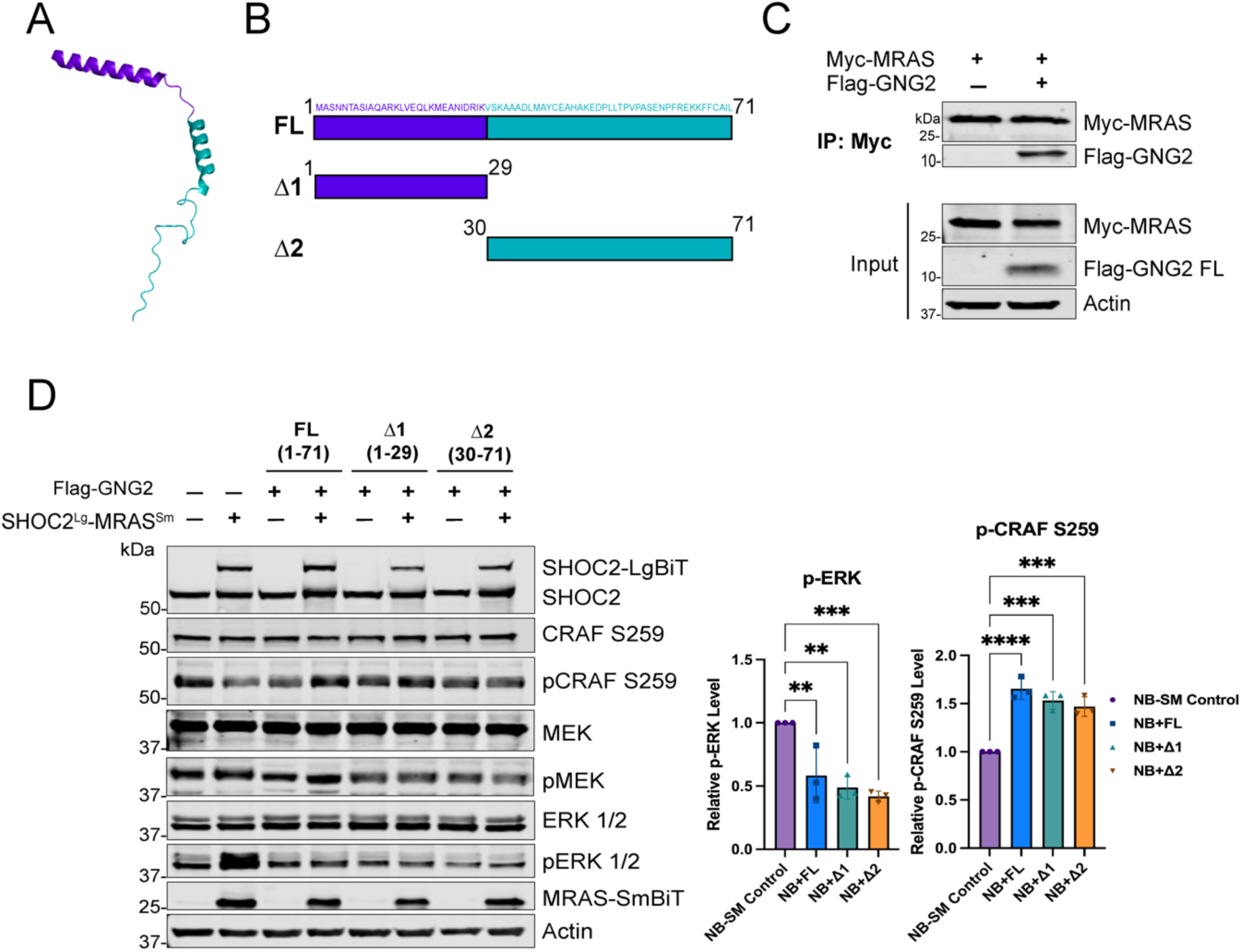
**GNG2 Variants Interact with MRAS and Inhibit MAPK Signaling**. **A**. AlphaFold2-predicted structure of FL GNG2 (residues 1–71), highlighting the sequences corresponding to the two truncated variants Δ1 (purple, residues 1-29) and Δ2 (cyan, residues 30-71). B. Schematic representation of FL GNG2 and its two truncated variants, Δ1 and Δ2. **C**. Co-immunoprecipitation analysis of Myc-MRAS and Flag-GNG2 (FL) in HEK293 cells. **D**. Immunoblot analysis of MAPK pathway components in HEK293 cells co-transfected with SHOC2^Lg^–MRAS^Sm^ NanoBiT constructs and Flag-tagged GNG2 FL, Δ1 and Δ2 variants at a 1:1 plasmid ratio. Quantification of phosphorylated ERK (p-ERK) and phosphorylated CRAF S259 (p-CRAF S259) levels from the immunoblots is shown alongside the corresponding blots. Data are representative of three biological replicates (n = 3). Statistical significance was determined by one-way ANOVA relative to the control (**P ≤ 0.01, ***P ≤ 0.001, ****P ≤ 0.0001).

We next examined whether GNG2 and its truncated variants affect MAPK signaling downstream of MRAS and SHOC2. HEK293 cells were co-transfected with the SHOC2^LgBiT^–MRAS^SmBiT^ NanoBiT construct and Flag-tagged GNG2 FL, Δ1, or Δ2 at an equimolar ratio. Immunoblot analysis showed that co-overexpression of SHOC2 and MRAS enhanced MAPK pathway signaling, as evidenced by increased MEK and ERK phosphorylation (p-MEK and p-ERK). In contrast, all three GNG2 constructs attenuated pathway activation, with GNG2 Δ2 exhibiting the strongest inhibition (∼60%), followed by GNG2 Δ1 (∼50%) and GNG2 FL (∼40%) (**Figure 1D**). These results indicate that GNG2 and its truncated variants suppress SHOC2/MRAS-dependent MAPK activation. Because SMP complex activity promotes RAF activation by dephosphorylating conserved inhibitory phosphoserine residues, we next examined the phosphorylation state of CRAF Ser259, a regulatory site associated with 14-3-3 binding and maintenance of RAF in an autoinhibited state (30–32). Expression of GNG2 increased CRAF Ser259 phosphorylation in cells overexpressing SHOC2 and MRAS, consistent with impaired RAF dephosphorylation (**Figure 1D**). Together, these findings suggest that GNG2 inhibits MAPK signaling, at least in part, by interfering with SMP complex function and thereby maintaining RAF in an autoinhibited state.

### GNG2 and Its Truncated Variants Disrupt SHOC2–MRAS Interactions and SMP Complex Assembly

To determine whether GNG2 binding to MRAS disrupts the SHOC2–MRAS interactions, we developed a live-cell NanoBiT bioluminescence assay in which SHOC2 and MRAS were fused to LgBiT and SmBiT, respectively, enabling real-time monitoring of SHOC2–MRAS interaction (**Figure 2A**). HEK293 cells expressing SHOC2^LgBiT^–MRAS^SmBiT^ were co-transfected with Flag-tagged full-length GNG2, Δ1, or Δ2. Expression of GNG2 and its variants significantly reduced NanoBiT luminescence by more than 75% compared with the control at an equimolar plasmid ratio (**Figure 2B**). Expression levels of SHOC2 and MRAS were constant across various conditions (**Figure 2B**); thus, the reduced NanoBiT luminescence primarily reflected disruption of the SHOC2– MRAS interaction rather than changes in protein abundance. Notably, the two truncated variants produced greater inhibition than FL GNG2, suggesting that the truncated proteins retain MRAS-binding determinants that might be less accessible in the full-length protein.

**Figure 2.**
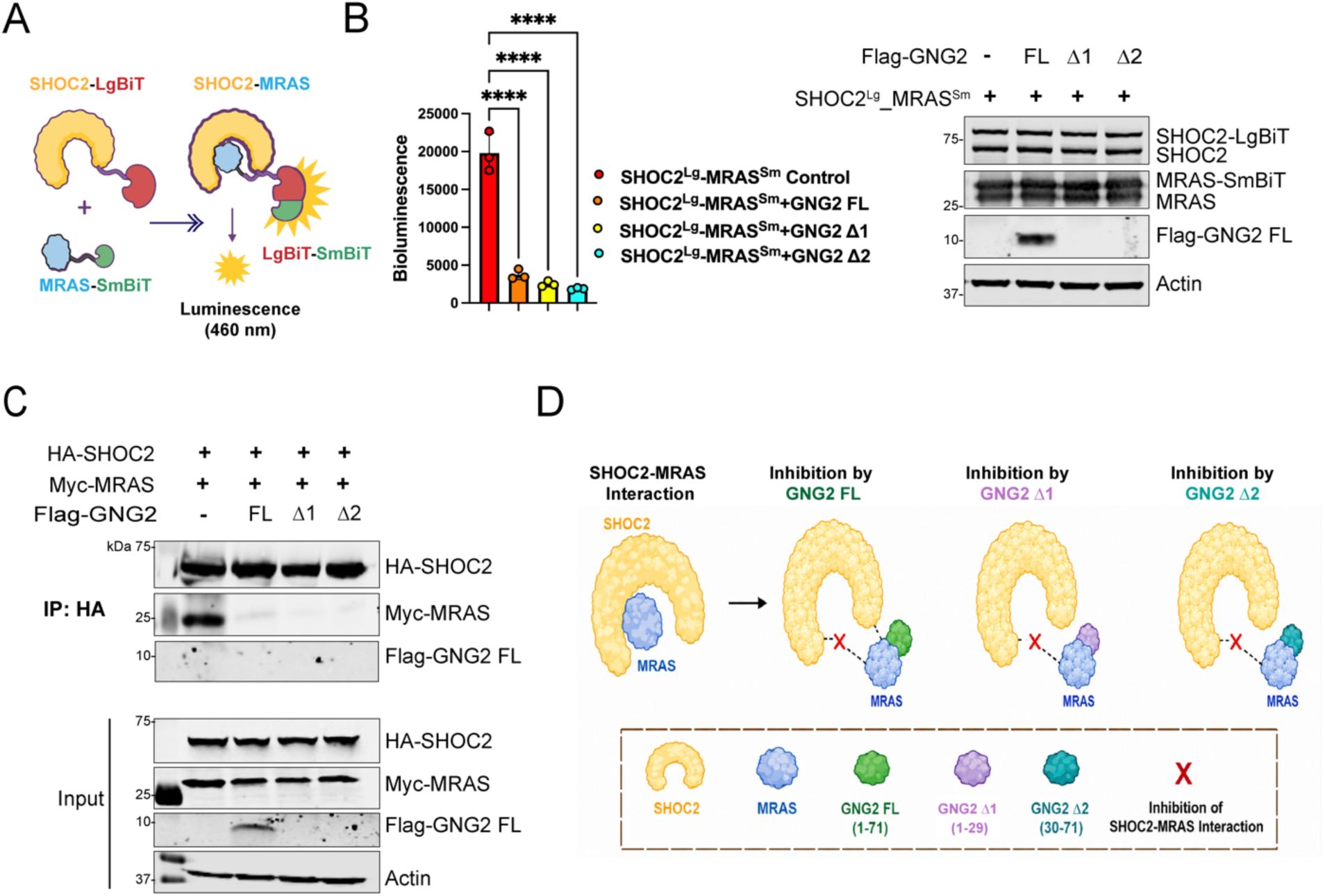
GNG2 and Its Truncated Variants Interact with MRAS and Disrupt SMP Complex Formation. **A**. Schematic representation of the NanoBiT assay used to monitor the SHOC2–MRAS interaction in live cells. Interaction between SHOC2– LgBiT and MRAS–SmBiT reconstitutes NanoLuc luciferase, producing a luminescent signal upon addition of the substrate (Furimazine). **B**. Live-cell NanoBiT assay of the SHOC2–MRAS interaction and corresponding immunoblot analysis of SHOC2^Lg^–MRAS^Sm^ and Flag-GNG2 expression in HEK293 cells co-transfected with SHOC2^Lg^–MRAS^Sm^ and Flag-GNG2 FL or the truncated variants (Δ1 and Δ2) at an equal (1:1) molar ratio. Bioluminescence signals were measured 48 h after transfection to assess the effect of GNG2 overexpression on the SHOC2–MRAS interaction. Data are presented as mean ± S.D. (n = 3 biological replicates). Statistical significance was determined using one-way ANOVA comparing NanoBiT control with GNG2-expressing samples (****P ≤ 0.0001). The Flag-GNG2-Δ1 and Δ2 were not readily detectable in the anti-Flag immunoblot, likely due to their small molecular sizes. **C**. Co-immunoprecipitation analysis of HA-SHOC2, Myc-MRAS, and Flag-GNG2 variants (FL/Δ1/Δ2). The Flag-GNG2-Δ1 and Δ2 were not readily detectable in the anti-Flag immunoblot, likely due to their small molecular sizes. **D**. Proposed model of GNG2-mediated inhibition of the SHOC2–MRAS interaction. Full-length GNG2 (FL, 1–71) and the truncated variants Δ1 (1–29) and Δ2 (30–71) bind MRAS, preventing its interaction with SHOC2 and thereby disrupting SHOC2–MRAS complex formation.

We next validated these findings using Co-IP. HEK293 cells were co-transfected with Myc-MRAS, HA-SHOC2, and Flag-tagged full-length GNG2, Δ1, or Δ2. Immunoprecipitation of HA-SHOC2 revealed a marked reduction in associated Myc-MRAS in the presence of all three GNG2 constructs (**Figure 2C**). GNG2 expression also reduced PP1Cα association with the SHOC2–MRAS complex (Figure S2). The truncated variants produced greater inhibition of SHOC2–MRAS association (>95%) than full-length GNG2 (>90%). In contrast, we did not detect a direct association between SHOC2 and full-length GNG2 under these conditions. These results are consistent with a model in which GNG2 interacts primarily with MRAS and interferes with the recruitment of SHOC2 and PP1Cα, rather than directly binding SHOC2. Because SHOC2–MRAS interaction is required for productive assembly of the SMP holophosphatase complex (17), these findings support a mechanism in which GNG2 suppresses MAPK signaling by disrupting SMP complex assembly upstream of RAF activation (**Figure 2D**).

### Molecular Dynamic Simulations Reveal GNG2 Binding to the Switch I/II Region of MRAS

To investigate the molecular basis of MRAS–GNG2 association, we performed all-atom MD simulations of MRAS in complex with either FL-GNG2 or the truncated Δ2, derived from GNG2 α-helical segments. The Δ1 fragment (residues 1–29) represents the N-terminal α-helix, while the Δ2 fragment (residues 30–71) represents the C-terminal α-helix and contains the CAAX motif required for membrane anchoring. Because Δ2 produced the strongest inhibition of MAPK signaling, we focused the MD analysis on full-length GNG2 and Δ2. Initial structures of the GNG2–MRAS complexes were generated using AlphaFold2-Multimer and Rosetta FlexPepDock. This integrative modeling independently yielded a consensus computational prediction localized at the Switch I (residues 40-48) and II (residues 70-86) regions of MRAS. In RAS-family GTPases, including KRAS, HRAS, and NRAS, Switch I undergoes nucleotide-dependent conformational changes and contributes to the primary interaction surface for downstream effectors, including RAF kinases (33). Switch II also undergoes conformational changes during the GTPase cycle and contributes to nucleotide hydrolysis and protein–protein interactions (34, 35). Both switch regions in the GTP analogue-loaded MRAS make direct contact with the leucine-rich repeat (LRR) domain of SHOC2, suggesting that these surfaces may contribute to competition between GNG2 and SMP complex components (15).

For the FL-GNG2–MRAS complex, MD simulations were performed with GNG2 tethered to a lipid bilayer composed of 80% POPC and 20% POPS (**Figure S3A**). The simulation quickly converged to a stable binding mode, showing persistent interactions between GNG2 and the Switch I/II region of GMPPNP-bound MRAS (**Figures 3A & 3B**). The residue contact map showed predominantly hydrophobic interactions between residues R13 and L15 in the N-terminal region of GNG2 with Y42, P44, L71, and F74 in MRAS. Additional contacts were observed involving residues GNG2 L37 and Y40 with MRAS Q80, L109, R83, and E79. A relatively stable salt bridge between GNG2 R13 and MRAS D41 further stabilized the interaction. The corresponding inter-residue distances remained within the ranges for persistent hydrophobic, salt-bridge, and hydrogen-bonding interactions throughout the trajectory (**Figure S4A**).

**Figure 3.**
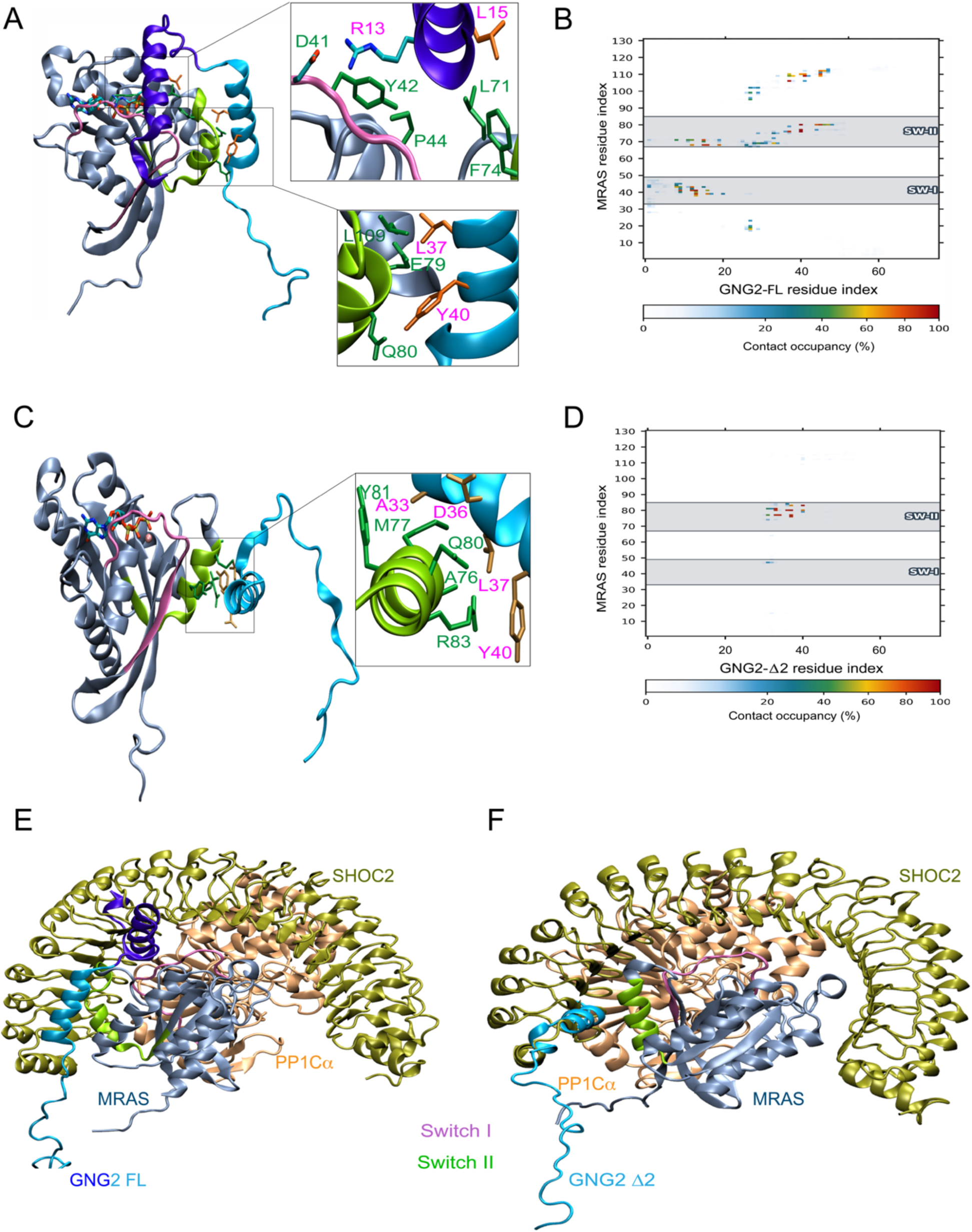
**Molecular Dynamic Simulation Analyses Reveal GNG2 Binding to the MRAS Switch I/II Interface and Competitively Blocks SHOC2 and PP1Cα Binding**. **A**. Representative molecular dynamics (MD) simulation snapshot showing GNG2 full-length (GNG2-FL; purple-cyan) bound to the Switch I (mauve) and Switch II (lime) regions of MRAS (ice blue; PDB: 7TVF). Interface residues are shown as licorice, and GTP and Mg²⁺ are displayed in licorice representation. **B**. Residue contact heat map of the MRAS–GNG2-FL complex. **C**. Representative MD simulation snapshot of the GNG2-Δ2 peptide (cyan) bound to the Switch II regions of MRAS. **D**. Residue contact heat map of the MRAS–GNG2-Δ2 complex. Contact maps were generated using a 5.0 Å heavy atom–heavy atom distance cut-off and demonstrate preferential peptide interactions with the Switch I and Switch II regions of MRAS. **E**–**F** AlphaFold2 Multimer models of GNG2-FL and GNG2-Δ2 bound to MRAS, superimposed onto the SHOC2–MRAS–PP1Cα (SMP) complex structure (PDB: 7TVF). The overlap shows that GNG2 occupies the SHOC2-binding surface on MRAS, supporting a competitive mechanism that disrupts SMP complex formation.

We next examined the GNG2 Δ2–MRAS complex also under membrane-tethered conditions. The Δ2 peptide retains the C-terminal CAAX motif and was therefore modeled with a geranylgeranyl lipid tail anchored to the membrane (**Figure 3C; Figure S3B**), as in the simulation of the FL-GNG2– MRAS complex. To better mimic the physiological conditions, MRAS was also modeled with its membrane-tethering farnesyl group. Similar to the full-length GNG2–MRAS simulation, the Δ2 peptide formed a stable interaction with the Switch II region of MRAS. The dominant contacts involved GNG2 A33, D36, L37, Y40 and MRAS A76, M77, Q80, R83, and Y81 (**Figure 3C** & **3D**; **Figure S4B**).

A replicate MD simulation confirmed that GNG2 FL and the Δ2 peptide interact with the Switch I/II interface of MRAS, as supported by residue-contact maps and stable interfacial interactions (**Figure S5**). These findings further support the proposed mechanism by which GNG2 interferes with SHOC2 binding to MRAS. Additional all-atom MD simulations with GDP-bound MRAS showed that GNG2 maintains stable interactions with the Switch I/II regions of MRAS (**Figure S6A**). Contact and distance analyses further supported these interactions (**Figures S6B & S6C**), suggesting that GNG2 can engage both GDP- and GTP-bound MRAS conformations, although the functional consequences of these interactions may depend on the nucleotide state.

Overall, our MD simulations revealed stable interactions between GNG2 and the Switch I and/or Switch II regions of MRAS. Importantly, these regions overlap with surfaces involved in SHOC2 and PP1Cα recognition within the SMP complex. MRAS residues F74, M77, Q80, and Y81 contribute to SHOC2 recognition, whereas D48, S49, L51, H53, and R83 participate in PP1Cα association (17). Within the SMP complex, MRAS H53 forms a key interaction with PP1Cα D179, and the H53A mutation reduces ternary complex formation by approximately 10-fold (17). Additionally, MRAS residues D48 and S49 in the Switch I region contact PP1Cα R188, a structural linchpin of the SMP complex that simultaneously engages MRAS and SHOC2 (17). These findings suggest that GNG2 may also interfere with PP1Cα engagement by occupying residues involved in the MRAS–PP1Cα interface, thereby hindering PP1Cα recruitment. Regardless, the assembly of SMP is contingent on each other, any disruption in the interface of protein–protein interactions leads to dissociation of the ternary complex, indicating multiple strategies can be applied to target this complex. This proposed mechanism provides a structural explanation for the observed inhibition of SMP complex formation and downstream MAPK signaling (**Figure 3E**&3**F**).

### GNG2 Δ2-Derived Peptides Inhibit SMP Complex Assembly and MAPK Signaling

Because the Δ2 fragment showed the strongest inhibition of MAPK signaling and efficiently disrupted SHOC2–MRAS association, we next sought to define a minimal inhibitory sequence. Four shorter GNG2-derived peptides were generated from the Δ2 region: Δ3 (residues 30-50aa), Δ4 (residues 51-71aa), Δ5 (residues 40-60aa), and Δ6 (residues 40-71aa) (**Figure 4A**). To investigate the structural basis for GNG2-mediated disruption of SMP assembly, we used AlphaFold2-Multimer to model the interactions of GNG2 Δ3–Δ6 peptides with MRAS. The predicted peptide–MRAS complexes were subsequently superimposed onto the SMP complex structure (PDB: 7TVF) to map the peptide-binding surfaces relative to the SHOC2–MRAS interface (**Figure S7**). Although the four peptides adopted distinct conformations and engaged different regions of MRAS, their predicted binding sites were positioned at or near the SHOC2-contacting surface in the assembled SMP complex. The longer Δ6 peptide (40–71) extended across a larger portion of the MRAS surface, potentially providing additional contacts and greater steric interference with SHOC2 engagement. Collectively, these structural models suggest that GNG2-derived peptides may disrupt SMP assembly by occupying or perturbing the MRAS surface required for productive SHOC2 association, providing a structural rationale for the experimentally observed inhibition of the SHOC2–MRAS interaction.

**Figure 4.**
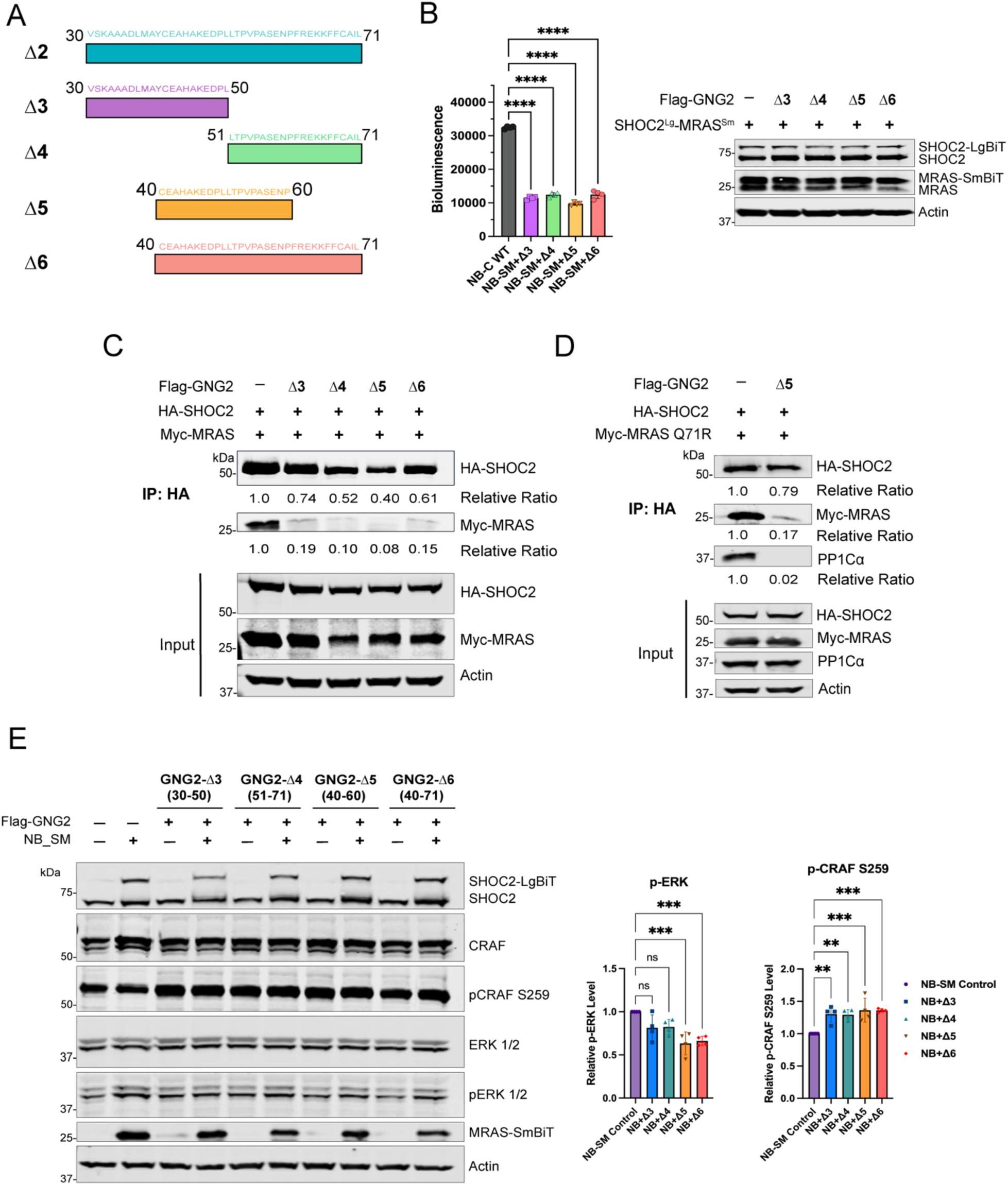
**GNG2 Δ2-derived truncated variants bind MRAS and suppress MAPK signaling**. **A**. Schematic representation of GNG2 Δ2-derived truncated variants: Δ3 (30–50), Δ4 (51–71), Δ5 (40–60), and Δ6 (40–71). **B**. Live-cell NanoBiT assay of the SHOC2–MRAS interaction and corresponding Immunoblot analysis of SHOC2^Lg^–MRAS^Sm^ expression in HEK293 cells co-transfected with SHOC2^Lg^–MRAS^Sm^ and Flag-tagged GNG2 variants (Δ3–Δ6) at plasmid ratios 1:1. Bioluminescence was measured 48h post-transfection to evaluate disruption of the SHOC2–MRAS interaction following GNG2 overexpression. Data are presented as mean ± S.D. (n = 4 biological replicates). Statistical significance was determined by one-way ANOVA relative to the NanoBiT control (***P ≤ 0.001, ****P ≤ 0.0001). **C**. Co-immunoprecipitation analysis of the interaction between HA-SHOC2 and Myc-MRAS WT in the presence of Flag-tagged GNG2 Δ3–Δ6 variants (n = 2 biological replicates). **D**. Co-immunoprecipitation analysis of HA-SHOC2 binding to the Myc-MRAS Q71R mutant in the presence of the Flag-tagged GNG2 Δ5 variant. **E**. Immunoblot analysis of MAPK pathway components in HEK293 cells co-transfected with SHOC2^Lg^–MRAS^Sm^ NanoBiT constructs (NB_SM) and Flag-tagged GNG2 Δ3–Δ6 variants at a 1:1 plasmid ratio (n = 4 replicates). Quantification of phosphorylated ERK (p-ERK) and phosphorylated CRAF S259 (p-CRAF S259) levels from the immunoblots is shown alongside the corresponding blots (n = 4 replicates). Statistical significance was determined by one-way ANOVA relative to the control (**P ≤ 0.01, ***P ≤ 0.001).

We next evaluated their effects on SHOC2–MRAS association using the live-cell NanoBiT assay. GNG2 Δ5 produced the strongest inhibition of the NanoBiT signal (∼75%), followed by Δ6 (∼68%) and Δ3 (∼65%) (**Figure 4B; Figure S8**). Consistent with these results, Co-IP analysis showed that all four peptides reduced the association between MRAS and SHOC2, with Δ5 producing the strongest inhibition (>80%) (**Figure 4C**).

We further tested whether GNG2-derived peptides could interfere with SHOC2 binding to the RASopathy-associated MRAS Q71R variant, which exhibits enhanced interaction with SHOC2. GNG2-derived peptide Δ5 reduced the SHOC2–MRAS Q71R–PP1Cα interaction, indicating that peptide-mediated inhibition is retained in this disease-associated MRAS background (**Figure 4D**). Consistent with their effects on SMP complex assembly, peptide expression increased CRAF Ser259 phosphorylation and reduced ERK phosphorylation by more than 45% (**Figure 4E**). These findings identify the GNG2 Δ2-derived peptides, particularly Δ5, as minimal sequences capable of disrupting SHOC2–MRAS association and suppressing downstream MAPK signaling.

### A Synthetic GNG2-Derived Peptide Directly Binds MRAS and Inhibits SMP Complex Assembly

To determine whether the inhibitory activity of the GNG2 Δ5 peptide results from direct interaction with MRAS, we synthesized a cell-permeable version of the peptide containing an N-terminal TAT sequence. Recombinant His-Flag-SHOC2, GST-MRAS, and GST-PP1Cα proteins were purified as described in the Methods (**Figure 5A**). BLI measurements demonstrated direct binding of the GNG2 Δ5 peptide to GMPPNP-bound MRAS (**Figure 5B**), exhibiting a dissociation constant (*K*_D_) of 257 ± 30 nM (*k*_on_ = 3.09 × 10^5^ M^-1^s^-1^ and *k*_off_ = 7.92 × 10^-2^ s^-1^). Surprisingly, the GNG2 Δ5 peptide exhibited comparably affinity for GDP-bound MRAS with *K*_D_ of 253 ± 77 nM (*k*_on_ = 1.95 × 10^5^ M^-1^s^-1^ and *k*_off_ = 4.93 × 10^-2^ s^-1^) (**Figure S9**), indicating that nucleotide loading has minimal impact on the measured binding affinity. Together with the MD simulation results, we propose that GNG2 recognition is not strictly dependent on the nucleotide state of MRAS. We next asked whether binding of the GNG2 Δ5 peptide to MRAS interferes with SMP complex assembly. GST-MRAS was pre-incubated with GNG2 Δ5 peptide for 2 h before the addition of GST-PP1Cα, and the resulting complex was analyzed for binding to immobilized SHOC2 by BLI. In the absence of peptide, the MRAS–PP1Cα complex displayed high affinity with SHOC2, with a *K*_D_ of 9 ± 4 nM (*k*_on_ = 1.25 × 10^4^ M^-1^s^-1^ and *k*_off_ = 1.12 × 10^-4^ s^-1^). Pre-incubation of MRAS with GNG2 Δ5 peptide markedly impaired SHOC2 association, increasing the apparent *K*_D_ to 1.88 ± 0.09 μM and resulting in a >200-fold reduction in apparent binding affinity (*k*_on_ = 2.96 × 10^3^ M^-1^s^-1^ and *k*_off_ = 5.54 × 10^-3^) (**Figure 5C&5D**). These findings demonstrate that the GNG2 Δ5 peptide directly binds MRAS and effectively disrupts assembly of the SMP complex.

**Figure 5.**
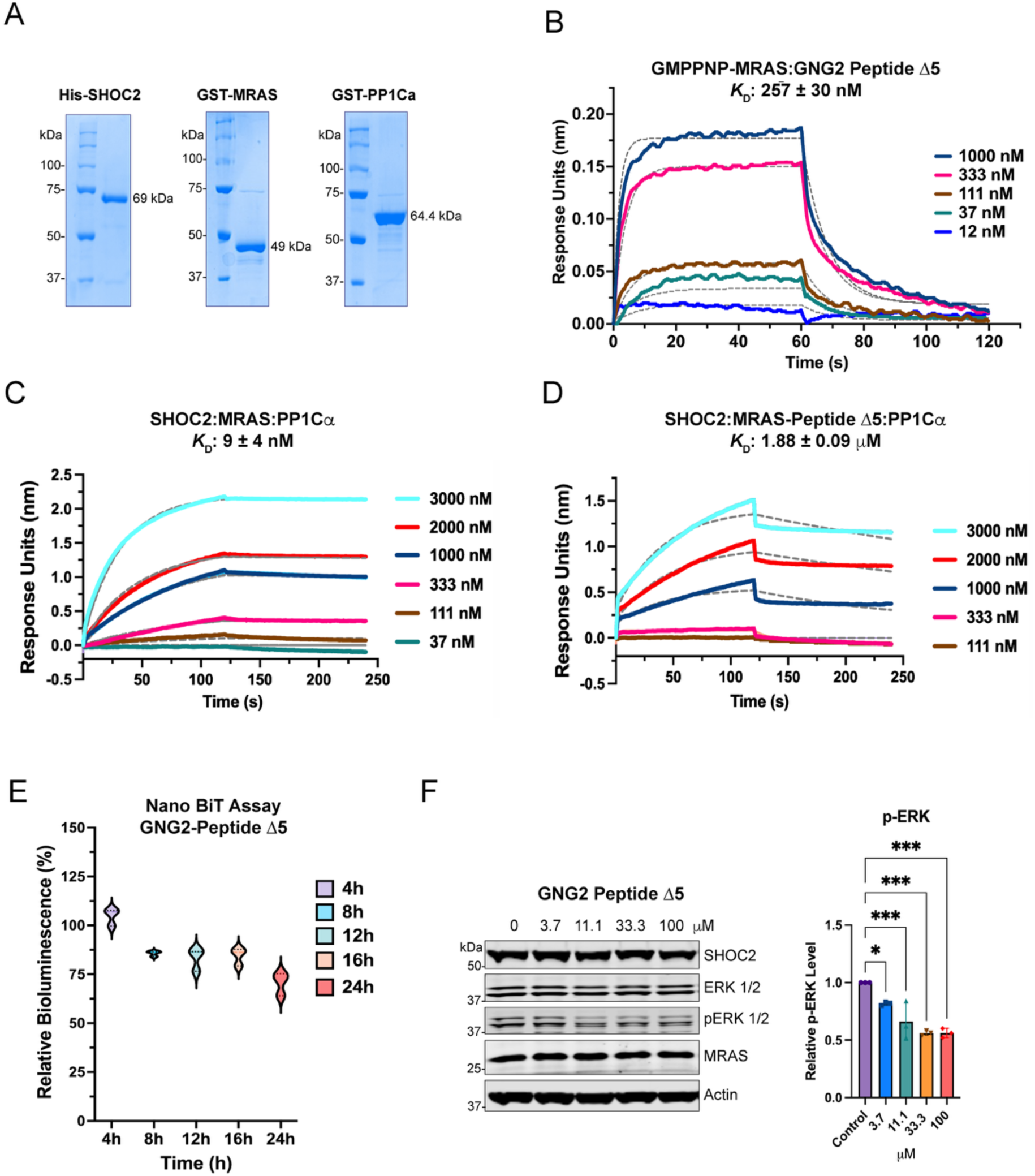
**TAT-conjugated GNG2 Δ5 peptide binds MRAS and disrupts SMP complex assembly**. **A**. Coomassie-stained SDS-PAGE analysis of purified recombinant His-SHOC2, GST-MRAS, and GST-PP1Cα proteins used for in vitro binding studies. **B**. Biolayer interferometry (BLI) analysis of the interaction between GPMPNP-MRAS and GNG2 Δ5 peptide. GST-MRAS (GMPPNP) was immobilized on GST biosensors, and increasing concentrations of GNG2 Δ5 peptide were used as analytes to determine binding kinetics. **C**. BLI analysis of SHOC2–MRAS–PP1Cα complex formation. His-SHOC2 was immobilized on His biosensors, and equimolar GMPPNP-MRAS and PP1Cα proteins were used as analytes. **D**. BLI analysis of the effect of GNG2 Δ5 peptide on SMP complex assembly. His-SHOC2 was immobilized on His biosensors, and PP1Cα with pre-incubated GMPPNP-MRAS–GNG2 Δ5 peptide complex (1:25 molar ratio) was used as the analyte to assess disruption of SHOC2–MRAS–PP1Cα complex formation. **E**. NanoBiT analysis of SHOC2–MRAS interaction in HEK293 cells following treatment with TAT-GNG2 Δ5 peptide (30 µM). **F**. Immunoblot analysis of MAPK pathway components in HEK293 cells treated with 16 h of increasing concentrations (0–100 µM) of TAT-GNG2 Δ5 peptide (n = 3 biological replicates). Quantification of p-ERK levels is shown alongside the corresponding immunoblots. Statistical significance was determined by one-way ANOVA relative to the control (*P ≤ 0.1, ***P ≤ 0.001).

We next evaluated the activity of the GNG2-derived peptide to disrupt the SHOC2–MRAS interaction in living cells using the NanoBiT assay. HEK293 cells expressing SHOC2^Lg^–MRAS^Sm^ NanoBiT constructs were treated with 30 μM peptides for 24 h. Luminescence was measured at 4h intervals to monitor the SHOC2–MRAS interaction. Peptide treatment reduced the SHOC2–MRAS NanoBiT signal by approximately 40% after 24 h, consistent with disruption of the interaction in cells (**Figure 5E**).

Finally, we evaluated the peptide’s effect on downstream MAPK signaling. Treatment of HEK293 cells with GNG2 Δ5 peptide (0–100 μM) for 16 h resulted in a concentration-dependent reduction in ERK phosphorylation, with significant inhibition observed at 33 and 100 μM peptide (**Figure 5F**). Together, these biochemical and cellular data provide proof of concept that a short GNG2-derived peptide can directly engage MRAS, interfere with SMP complex assembly, and suppress downstream MAPK signaling.

## Discussion

The structural and functional characterization of the SHOC2–MRAS–PP1Cα (SMP) holophosphatase complex has identified an important regulatory checkpoint upstream of RAF activation. By dephosphorylating the conserved inhibitory serine residues within RAF (ARAF Ser214, BRAF Ser365, and CRAF Ser259), the SMP complex promotes release of RAF autoinhibition and enables RAF dimerization and downstream MEK–ERK signaling (15–18).

Because productive SMP activity depends on assembly of a ternary complex, disruption of any of its essential PPIs can potentially suppress RAF activation. Our findings identify GNG2 as a regulator of this process and provide evidence that GNG2 suppresses MAPK signaling by competing with SHOC2 for interaction with MRAS.

The central finding of this study is that GNG2 directly engages MRAS and interferes with the SHOC2–MRAS association. NanoBiT and Co-IP experiments demonstrated that full-length GNG2 and its truncated variants markedly reduce SHOC2–MRAS interaction in cells, whereas GNG2 itself was not detected in association with SHOC2. These findings support a model in which GNG2 acts primarily through MRAS rather than through direct interaction with SHOC2. The functional consequences are inhibition of SMP complex assembly, increased phosphorylation at the inhibitory CRAF Ser259 site, and attenuation of MEK and ERK activation. Structural studies of the Gβ1γ2 complex (PDB entries 1GOT (36) and 1OMW (37)) show that GNG2 interacts with Gβ1 through its N-terminal helix and the β-propeller domain. Gβ1 association may influence GNG2 conformation or accessibility for MRAS binding, potentially modulating SHOC2–MRAS interaction and SMP assembly. However, the role of Gβ1 in GNG2-mediated inhibition of SMP formation remains to be investigated. Overall, our data expand the known functions of GNG2 beyond its canonical role as a component of the heterotrimeric G-protein complex and suggest that GNG2 can influence MAPK signaling through direct regulation of an MRAS-dependent signaling complex.

Our MD simulations provide a structural rationale for this proposed mechanism. Full-length GNG2 and the Δ2 fragment formed stable interactions with the Switch I/II region of MRAS, and the predicted binding surfaces overlap with regions required for SHOC2 and PP1Cα recognition. In particular, MRAS residues F74, M77, Q80, and Y81 contribute to SHOC2 binding, while additional residues within the Switch I/II region participate in PP1Cα recruitment (15, 17, 38). The GNG2-binding mode identified in our simulations therefore places GNG2 on a strategically important MRAS surface that could interfere with the assembly of the SMP complex. Although direct structural determination of the GNG2–MRAS complex will be required to definitively establish the interaction, the convergence of AlphaFold-based modeling, Rosetta docking, molecular dynamics simulations, BLI, NanoBiT, and Co-IP provides strong support for a competitive mechanism centered on the MRAS Switch I/II interface. The ability of GNG2 to maintain predicted interactions with both GDP- and GMPPNP-bound MRAS suggests that its recognition of MRAS may not be strictly dependent on nucleotide state, raising the possibility that GNG2 interferes with SHOC2–MRAS–PP1Cα (SMP) complex assembly across distinct MRAS conformational states rather than solely by targeting the active-like conformation.

The identification of a minimal GNG2-derived inhibitory peptide further demonstrates that the MRAS-binding activity of GNG2 can be separated from the remainder of the protein. The Δ5 peptide directly bound to inactive (GDP) as well as active (GMPPNP) MRAS with a *K*_D_ of 256 nM in BLI experiments and reduced SHOC2 association with MRAS by more than 200-fold in the reconstituted system. Importantly, the peptide also reduced SHOC2–MRAS interaction and ERK phosphorylation in cells, establishing proof of concept that the MRAS-binding interface can be pharmacologically modulated using a short peptide. The activity of the peptide against the RASopathy-associated MRAS Q71R variant is particularly noteworthy because this mutation enhances SHOC2 interaction and promotes MAPK pathway activation. These findings raise the possibility that direct modulation of the MRAS–SHOC2 interface could be useful not only in RAS-driven malignancies but also in pathological conditions driven by hyperactive SMP signaling.

The therapeutic implications of targeting the SMP complex are also noteworthy. Unlike catalytic-site inhibitors of MEK or ERK, which act downstream of RAF activation, or RAS-directed inhibitors that alter RAS nucleotide state, GNG2-derived peptides target a protein–protein interaction required for productive SMP assembly. This mechanism may provide an alternative means of suppressing RAF activation and could potentially be combined with existing MAPK pathway inhibitors. However, the present peptide remains a proof-of-concept molecule rather than a therapeutic candidate. Although GNG2 Δ5 binds MRAS with submicromolar affinity in vitro, higher concentrations were required to suppress cellular ERK phosphorylation, consistent with limitations associated with intracellular peptide delivery, stability, and competition with endogenous SMP complexes. Optimization of the peptide through sequence engineering, stapling, or cyclization, and improved intracellular delivery may therefore be necessary to achieve more potent cellular activity.

In summary, our findings identify GNG2 as a previously unrecognized regulator of SHOC2-dependent RAF activation. GNG2 directly engages MRAS through a surface overlapping the SHOC2-recognition region, thereby interfering with SMP complex assembly and maintaining RAF in its inhibitory phosphorylated state. Importantly, this endogenous regulatory mechanism can be reduced to a short GNG2-derived peptide that directly binds MRAS and suppresses SMP-dependent MAPK signaling in vitro and in cells. Together, these findings establish the MRAS–SHOC2 interface as a tractable protein–protein interaction for regulating RAF activation and provide a molecular framework for developing GNG2-derived peptide modulators of RAS–RAF–MAPK signaling.

## Experimental Section

### Plasmids

The bacterial expression constructs GST-MRAS (plasmid #109934), GST-PP1Cα (plasmid #55661), and mammalian mApple-GNG2 (plasmid #133857) were obtained from Addgene. Human MRAS (Addgene plasmid #70419) was cloned into the Myc-pcDNA3.1 vector, whereas wild-type SHOC2 (Addgene plasmid #190568) was cloned into the HA-pRc vector for mammalian expression and into pFastBac for baculovirus-mediated expression in Sf9 insect cells using Gibson Assembly (New England Biolabs, NEB). The mApple tag of GNG2 was replaced with the Flag tag using the KLD reaction (NEB). Further truncated variants of Flag-GNG2 (Δ1-Δ6) were made using Flag-GNG2 FL (1–71) as a template in the KLD reaction. For a live-cell bioluminescence assay, SHOC2 and MRAS were cloned with the N-terminus LgBiT and SmBiT of the NanoBiT plasmid, respectively, using Gibson assembly. Plasmid prep was performed according to the manufacturer’s protocol (Zymo Research D4201).

The following primers were used:

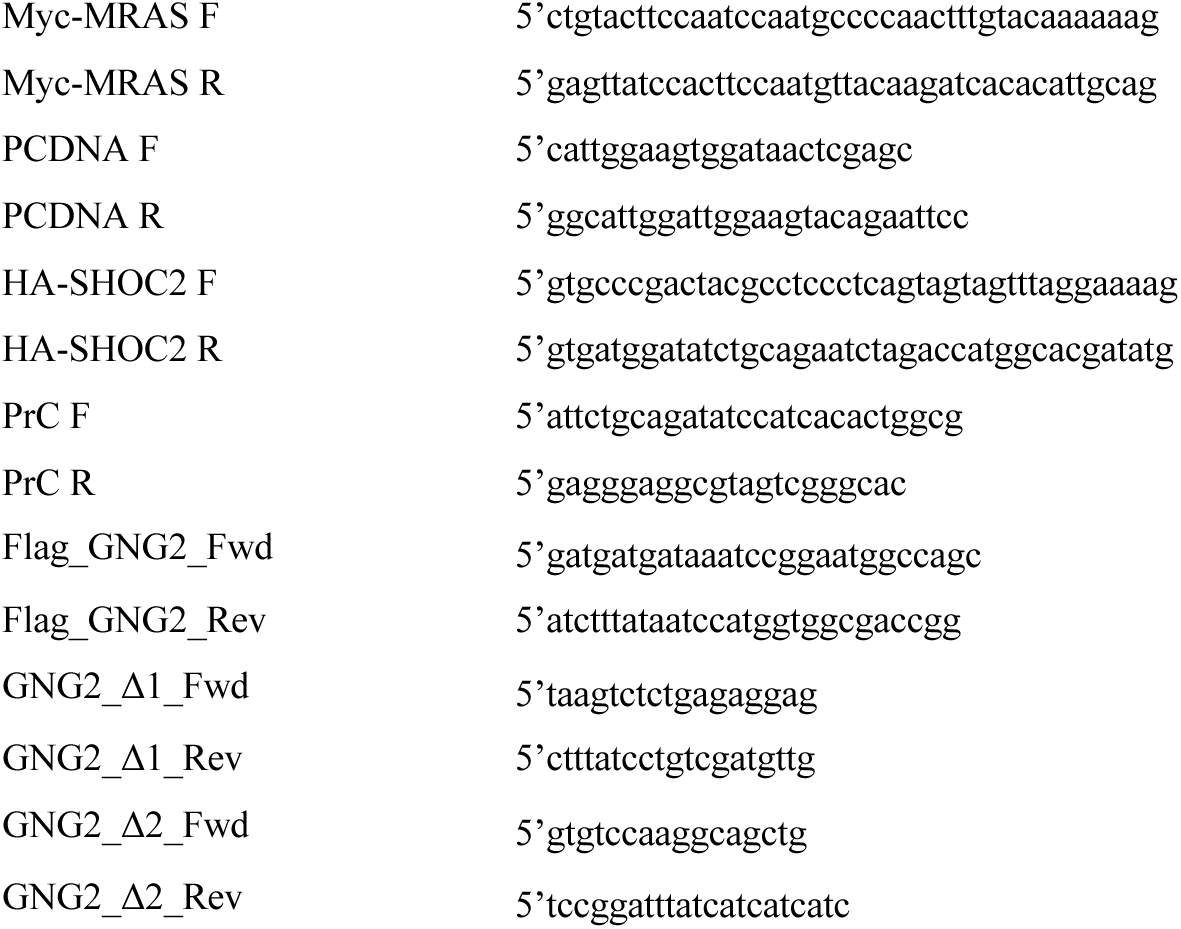

### Bacterial Protein Expression

The plasmid constructs (GST-MRAS and GST-PP1Cα) were transformed into BL21 codon+ *E. coli* cells and grown to an OD_600nm_ 0.6−0.8 in LB broth (PP1Cα was supplemented with 1 mM MnCl_2_), followed by induction with 0.5 mM IPTG. Cells were left overnight at 18 °C, shaking at 210 rpm. Cells were pelleted, flash-frozen, and stored at −80 °C.

### Protein Purification SHOC2 Purification

Full-length SHOC2 was cloned into the pFastBac vector to express an N-terminally tagged 6×His-Flag-SHOC2 fusion protein. Recombinant baculovirus was generated using the corresponding bacmid and used to infect Sf9 insect cells. Cells were harvested by centrifugation 72 h post-infection and resuspended in lysis buffer containing 50 mM HEPES (pH 7.5), 500 mM NaCl, 10% glycerol, 0.5 mM TCEP, 0.001% NP-40, and protease inhibitor cocktail. Cells were lysed using a Dounce homogenizer (50 strokes), and cell debris was removed by high-speed centrifugation. The clarified lysate was supplemented with 20 mM imidazole and incubated with pre-equilibrated cobalt affinity resin for 2h at 4 °C. After washing with buffer containing 50 mM HEPES (pH 7.5), 500 mM NaCl, 10% glycerol, and 0.5 mM TCEP, the bound protein was eluted using a linear imidazole gradient (90–400 mM). Fractions containing SHOC2 were pooled and further purified by size-exclusion chromatography on a Superdex 200 Increase 10/300 GL column (Cytiva) equilibrated with 50 mM HEPES (pH 7.4), 150 mM NaCl, and 10% glycerol. Protein purity was assessed by SDS-PAGE followed by Coomassie Brilliant Blue staining. Purified protein was concentrated, quantified using a BSA standard curve, aliquoted, flash-frozen in liquid nitrogen, and stored at −80 °C.

### MRAS Purification

The cell pellet obtained from a 500 mL bacterial culture was resuspended in 20 mL of lysis buffer (50 mM Tris-HCl, pH 8.0, 500 mM NaCl, 0.5 mM MgCl_2_, 10% glycerol, and 0.5 mM TCEP) supplemented with protease inhibitor cocktail. The whole-cell lysate (35 mL) was lysed by six passes through a French pressure cell press at 1,720 psi and centrifuged at 18,000 rpm for 40 min at 4 °C. The clarified supernatant was incubated with 0.5 mL of pre-equilibrated glutathione resin for 2h at 4 °C. After extensive washing (five column volumes) with wash buffer (50 mM Tris-HCl, pH 8.0, 500 mM NaCl, 0.5 mM MgCl_2_, 10% glycerol, and 0.5 mM TCEP), bound proteins were eluted by centrifugation (500 × g, 90 s) using elution buffer containing 20 mM reduced glutathione.

For nucleotide exchange, the purified proteins were incubated in HEPES buffer containing 10 mM EDTA, and a 10-fold molar excess of the non-hydrolyzable GTP analog GMPPNP (MilliporeSigma) for 30 min at 4 °C. MgCl_2_ was then added to a final concentration of 20 mM to promote nucleotide rebinding, and the mixture was incubated for an additional 1 h 30 minutes at 4 °C with gentle rotation. Proteins were further purified by size-exclusion chromatography on a Superdex 200 10/300 GL column (Cytiva) equilibrated with 50 mM HEPES (pH 7.4), 150 mM NaCl, and 10% glycerol. Protein purification was monitored by SDS-PAGE followed by Coomassie Brilliant Blue staining. Purified proteins were concentrated, quantified using a BSA standard curve, aliquoted, flash-frozen in liquid nitrogen, and stored at −80 °C until further use.

### PP1Cα Purification

GST-PP1Cα cell pellets were resuspended in 20 mL of lysis buffer containing 50 mM Tris-HCl (pH 8.0), 500 mM NaCl, 0.5 mM MnCl₂, 10% glycerol, and 0.5 mM TCEP, supplemented with protease inhibitor cocktail. The cell suspension (up to 35 mL) was lysed using a French pressure cell press at 1,520 psi and clarified by centrifugation at 18,000 rpm for 40 min at 4 °C. The supernatant was incubated with pre-equilibrated glutathione resin for 2h at 4 °C. After extensive washing (five column volumes) with wash buffer (50 mM Tris-HCl, pH 8.0, 500 mM NaCl, 0.5 mM MnCl₂, 10% glycerol, and 0.5 mM TCEP), bound GST-PP1Cα was eluted with elution buffer containing 20 mM reduced glutathione. The eluted protein was further purified by size-exclusion chromatography on a Superdex 200 Increase 10/300 GL column (Cytiva) equilibrated with 50 mM HEPES (pH 7.4), 150 mM NaCl, 0.5 mM MnCl₂, and 10% glycerol. Protein purity was monitored by SDS-PAGE followed by Coomassie Brilliant Blue staining. Purified protein fractions were concentrated, quantified using a BSA standard curve, aliquoted, flash-frozen in liquid nitrogen, and stored at −80 °C.

### Biolayer Interferometry (BLI)

Binding studies of GDP- or GMPPNP-bound MRAS- and TAT-tagged GNG2 peptide Δ5 were measured using an Octet N1 (Sartorius). GST-MRAS (15 ng) was immobilized on the GST-sensor, and GNG2 peptide 5 was used as the analyte (0-1000 nM) in the buffer (20 mM HEPES, pH 7.4, 150 mM NaCl, 0.05% glycerol) and allowed to associate for 60 s, followed by dissociation for 2 min. For binding studies of SHOC2 with MRAS and PP1Cα, 20 ng of His-SHOC2 was immobilized onto Ni- NTA (HSK) biosensors. GST-MRAS, pre-incubated in the presence or absence of GNG2 Δ5 peptide for 2 h, was mixed with GST-PP1Cα and used as the analyte at concentrations ranging from 0 to 3000 nM. All measurements were performed in an assay buffer containing 20 mM HEPES (pH 7.4), 150 mM NaCl, 0.05% Tween-20, and 1 mM TCEP. Association was monitored for 2 min, followed by a 2 min dissociation phase. All data were normalized and analyzed using the global-fit binding (1:1) model across all concentrations with the Octet Analysis Studio 12 software provided by Sartorius to derive K*_D_* values, and the graphs were generated using GraphPad Prism software.

### Cell Culture and Transfection

HEK293 cells were cultured at 37°C and 5% CO_2_ until 70-90% confluency, then passaged. New cell stocks were thawed after 12 passages. Cells were kept in Dulbecco’s Modified Eagle Medium (DMEM) supplemented with 1% Penicillin-Streptomycin, 1% L-Glutamate, and 10% Fetal Bovine Serum. Small-scale transfections of 1 million cells were plated in a 6-well plate and allowed to reach 60% confluency before transfection. After incubation for 24 h, 1-2 μg of plasmid DNA was added, along with 3-6 μg of PEI-MAX per well. Transfections were cultured for 48 h at 37°C, 5% CO_2_. The media was aspirated, and the cells were washed with cold PBS. Cells were resuspended in 1 mL of PBS (for further use), centrifuged, flash-frozen in liquid nitrogen, and then lysed with 4% SDS. Protein concentrations were determined by the BCA assay; 20-30 µg of protein was loaded onto an SDS-PAGE gel, and the gel was visualized by immunoblotting.

For Co-IP experiments, HEK293 cells were seeded at 5 million cells per 10-cm dish and transfected at 60% confluency with a 1:3 DNA: transfectant ratio (PEI-MAX). Single transfections were carried out with 10 μg in a 10-cm dish. The HA-SHOC2, Flag-GNG2 (FL/Δ1-6, and Myc-MRAS were carried using a 1:1:1 ratio. Transfections were cultured for 48 h at 37 °C in 5% CO_2_ before harvest in cold PBS. Cells were lysed using a modified radioimmunoprecipitation assay buffer (50 mM HEPES pH 7.4, 150 mM NaCl, 0.1% NP40, 1 mM EDTA, 5% glycerol, 1 mM PMSF, protease inhibitor (Roche 11836170001)) at 4 °C, with 2 h of rotation. Immunoprecipitation was carried out according to the manufacturer’s protocols for the immunoprecipitation of HA-tagged/Flag-tagged proteins (Thermofisher) and quenched in 4× Laemmli loading dye for 5 min at 95 °C.

The effect of synthesized TAT-tagged GNG2 Δ5 peptides on HEK293 cells: cells were treated with different peptide concentrations (0-100 μM) for 16 h in Opti-MEM supplemented with 1 % FBS, followed by peptide replenishment and an additional 4 h incubation before protein extraction.

### Immunoblot analysis

Samples were diluted in 4X loading dye and heated to 95 °C for 5 minutes. SDS PAGE gels were run at 90 V for 100 minutes. Primary antibodies (HA/V5/Myc/Flag/MEK/pMEK/ERK/pERK/CRAF/pCRAF-S259/SHOC2/MRAS/Actin) were incubated overnight with shaking at 4°C. The secondary antibody (IRDye, Licorbio) incubation was performed for 2 h at room temperature with shaking. Membranes were visualized by using a LICOR Odyssey CLx scanner. Quantification of p-MEK and p-ERK levels was performed using ImageJ software.

### Live-cells NanoBiT bioluminescence assay

HEK293 cells were cultured at 37°C with 5% CO_2_ until 70-90% confluency, then passaged. Cells were kept in DMEM supplemented with 1% Penicillin-Streptomycin, 1% L-Glutamate, and 10% Fetal Bovine Serum. Small-scale transfections of 1 million cells were plated in a 6-well plate and allowed to reach 60% confluency before transfection. After incubation for 24 h, 1.5 μg of SHOC2^Lg^– MRAS^Sm^ WT NB plasmid DNA, with or without Flag-GNG2 (FL/1-6), was added to 4.5 μg PEI-MAX (1:3) per well. After 24h incubation, cells were counted and reseeded with 50k-80k cells/well (100 µL) in Opti-DEM supplemented with 4% FBS into the 96-well plate and incubated for another 24 h at 37°C, 5% CO_2_. The substrate furimazine (10 μM) was added to all wells, and luminescence was measured after 5 minutes of incubation. The plate was further read by a microplate reader by light-emission acquisition at 485 and 568 nm. For western blot analysis, cells were resuspended in 1 mL PBS, immediately lysed with 4% SDS, and analyzed by western blot.

The effect of synthesized TAT-tagged GNG2 Δ5 peptides on the SHOC2–MRAS interaction was evaluated using the NanoBiT bioluminescence complementation assay in HEK293 cells. Cells were transfected with 2 μg of SHOC2^Lg^– MRAS^Sm^ WT NB plasmid. Following transfection, cells were allowed to express the NanoBiT constructs for 24 h. After 24h incubation, cells were counted and reseeded with 30k cells/well (100 µL) in Opti-DEM into the 96-well plate and incubated for another 24 h at 37°C, 5% CO_2_. The cells were treated with synthesized GNG2 Δ5 peptides at a final concentration of 30 μM. To maintain peptide availability throughout the experiment, fresh peptide was added every 4 h during the 24 h treatment period. NanoBiT luminescence was measured immediately using the furamazine substrate. The luminescence was measured after 5 minutes of incubation. The plate was further read by a microplate reader by light-emission acquisition at 460 nm and 568 nm. The changes in NanoBiT luminescence were used as a quantitative measure of the SHOC2–MRAS interaction.

### Molecular Dynamics Simulations

Simulations were carried out using the Amber25 software package, with the FF19SB and Lipids21 force fields for proteins and the lipid bilayer, respectively. MRAS’s initial structure was derived from the crystal structure of the SHOC2– MRAS– PP1CΑ complex PDB: 7TVF (17), while Alphafold (39) was used to generate the initial structure of FL GNG2. Δ1 or Δ2 represent the N-terminal and C-terminal segments, respectively. Initial structures of the protein-protein (MRAS-GNG2) complexes were generated using the AlphaFold2 multimer (36) and compared to Rosetta FlexPepDock (40) lowest REU conformations based on Interface scores (I_sc). The RESP-charge fitting method (41) and the General Amber force field (GAFF) (42) were applied to the farnesyl lipid tail for MRAS and the GNG2 geranylgeranyl lipid tail. The FL GNG2-MRAS and GNG2 Δ2-MRAS systems were placed together with a lipid bilayer (periodic in two dimensions) consisting of 80% POPC (1-Palmitoyl-20-oleoylphosphatidylcholine) and 20% POPS (1-Palmitoyl-2-oleoylphosphatidylserine) with the farnesyl and geranylgeranyl lipid moiety pointing towards the bilayer surface. Each protein-peptide or protein-peptide/lipid system was solvated in a periodic box with explicit OPC water molecules and appropriate counterions (Na^+^ or Cl^-^) to ensure an overall neutral charge of the system.

Equilibration of each system began with initial energy minimization to remove steric clashes. The systems were then gradually heated to 300 K, followed by equilibration of the simulation box size at constant pressure (NPT) at 1 atm and 300 K. Following equilibration, each system was simulated for a 500ns production run in the NVT ensemble at 300 K with a timestep of 1 fs. Structural analysis of the MD trajectories was performed using the Amber25 cpptraj program.

| **Antibody** | **Manufacturer** | **Catalog Number** |
|---|---|---|
| HA | Cell Signaling | 3724 |
| Myc | Cell Signaling | 2272 |
| MEK1/2 | Cell Signaling | 4694 |
| pMEK S217/S222 | Cell Signaling | 9154 |
| ERK1/2 | Cell Signaling | 4696 |
| pERK T202/Y204 | Cell Signaling | 4370 |
| SHOC2 | Cell Signaling | 53600 |
| β -Actin | Cell Signaling | 3700 |
| p-CRAF S259 | Cell Signaling | 9421s |
| MRAS | Antibodies.com | 77222-714 |
| GNG2 | AFG Bioscience | 7711-115 |
| Flag | Sigma-aldrich | F1804 |

