## Supplemental file for "GNG2–MRAS interaction regulates SHOC2-dependent RAF activation and MAPK signaling"

#### Supplementary Information

- Supplementary Figures 1-9

#### Figure S1

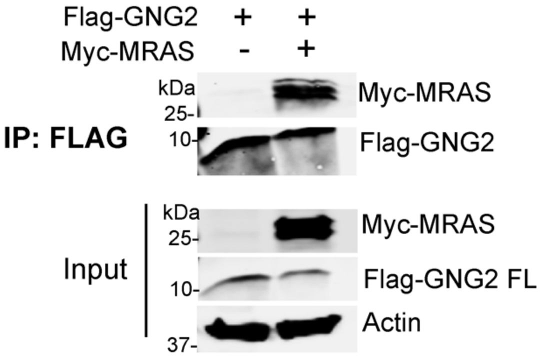


**FIGURE S1. Co-immunoprecipitation analysis of the GNG2–MRAS interaction.** Co-immunoprecipitation of Flag-GNG2 (full-length, FL) in HEK293 cells in the presence or absence of Myc-MRAS. Flag affinity resin was used to immunoprecipitate Flag-GNG2, and the precipitates were analyzed for the presence of Myc-MRAS.

**Figure S2**

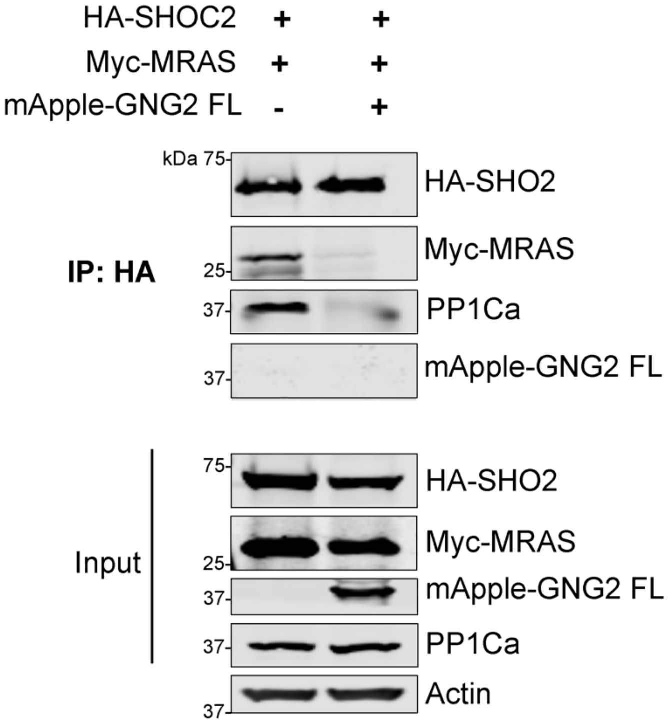


**FIGURE S2. Co-immunoprecipitation analysis of the GNG2–MRAS interaction.** Co-immunoprecipitation analysis of HA-SHOC2, Myc-MRAS, and mApple-GNG2 FL. HA affinity resin was used to immunoprecipitate HA-SHOC2, and the precipitates were analyzed for the presence of Myc-MRAS, PP1Ca (endogenous) and mApple-GNG2 FL.

### Figure S3

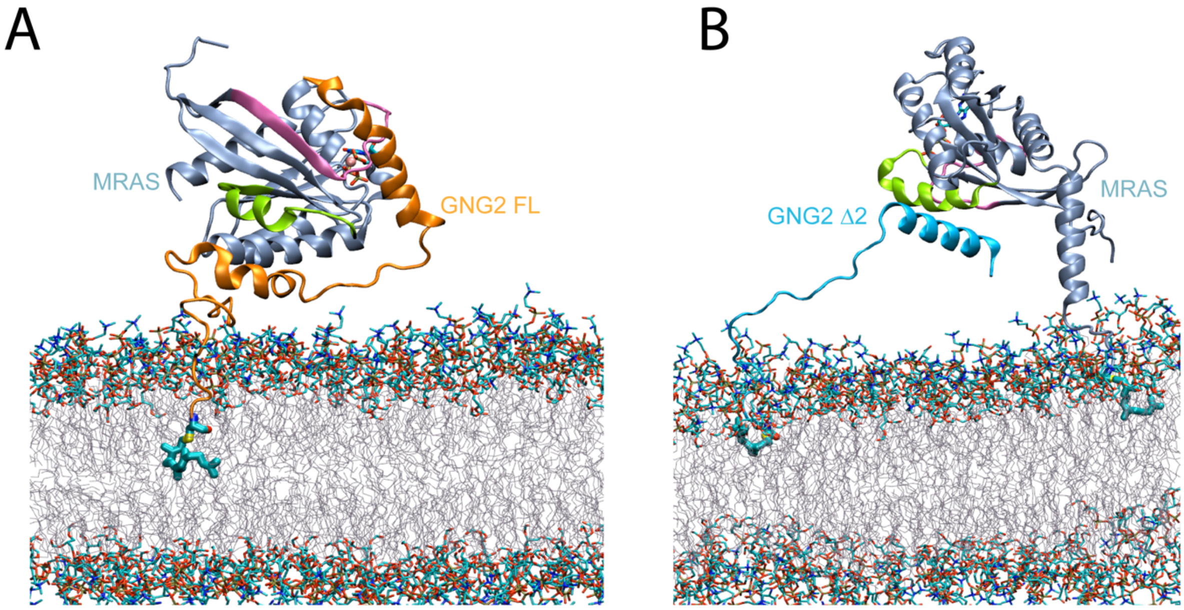


**FIGURE S3. Molecular dynamics models of GNG2 and MRAS at the membrane. A-B**. Initial molecular dynamics (MD) models of full-length GNG2 (orange) (**A**) and the GNG2-Δ2 peptide (teal) (**B**) in complex with MRAS (ice blue). Switch I and Switch II regions of MRAS are shown in mauve and lime, respectively. Lipid anchors (farnesyl and geranylgeranyl) are shown as cyan licorice representations. Membrane phospholipid headgroups, phosphotidylcholine (PC) and phosphatidylserine (PS) are shown in licorice representation, and the lipid acyl tails, palmitic acid (PA) and oleic acid (OL) are displayed as line representation.

### Figure S4

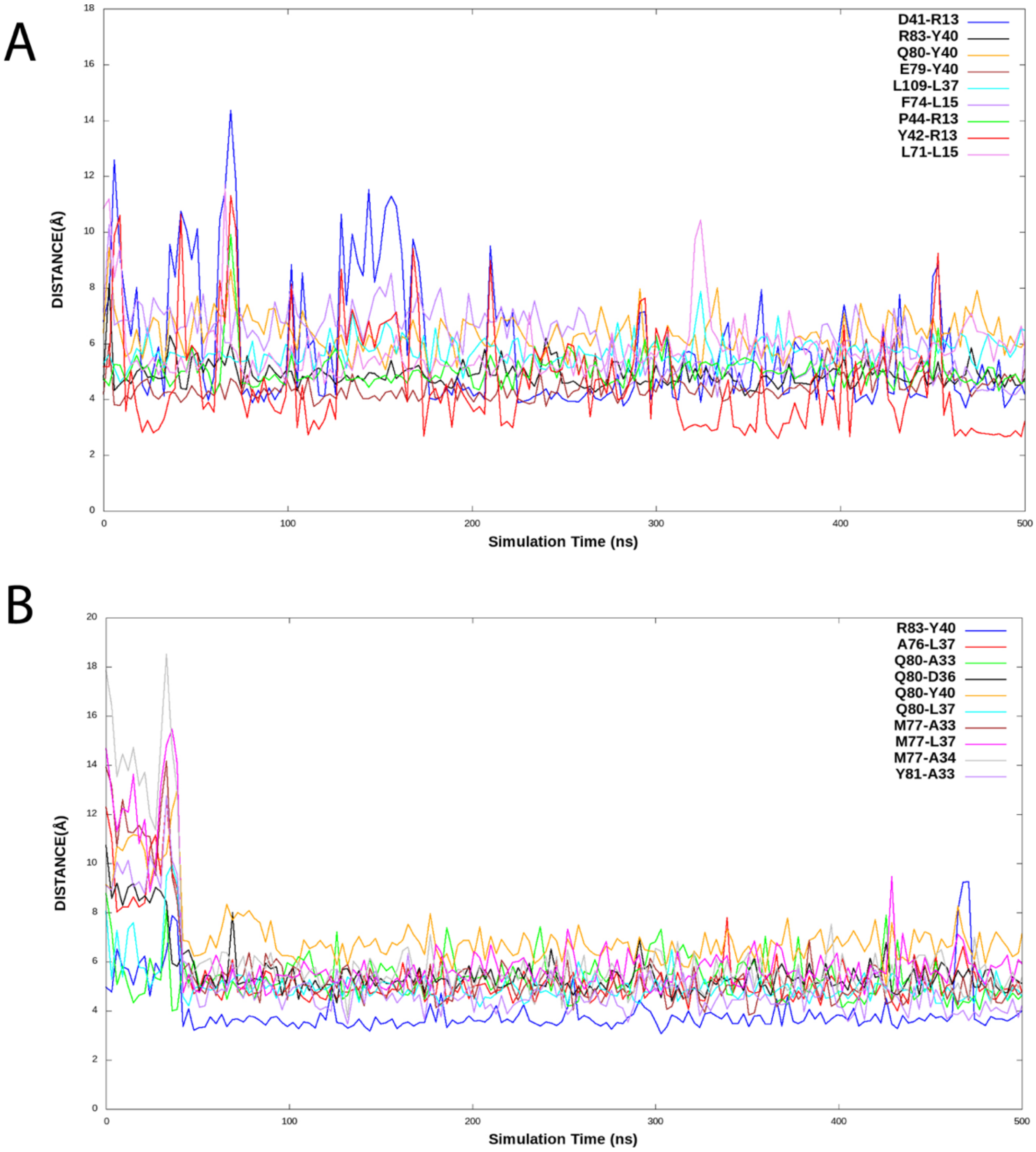


**FIGURE S4. Stability of GNG2–MRAS interactions during molecular dynamics simulations.** Distance plots demonstrating the stability of representative hydrophobic interactions, hydrogen bonds, and salt bridges between MRAS and (**A**) FL-GNG2 and (**B**) the GNG2-Δ2 peptide over the course of the molecular dynamics simulations.

### Figure S5

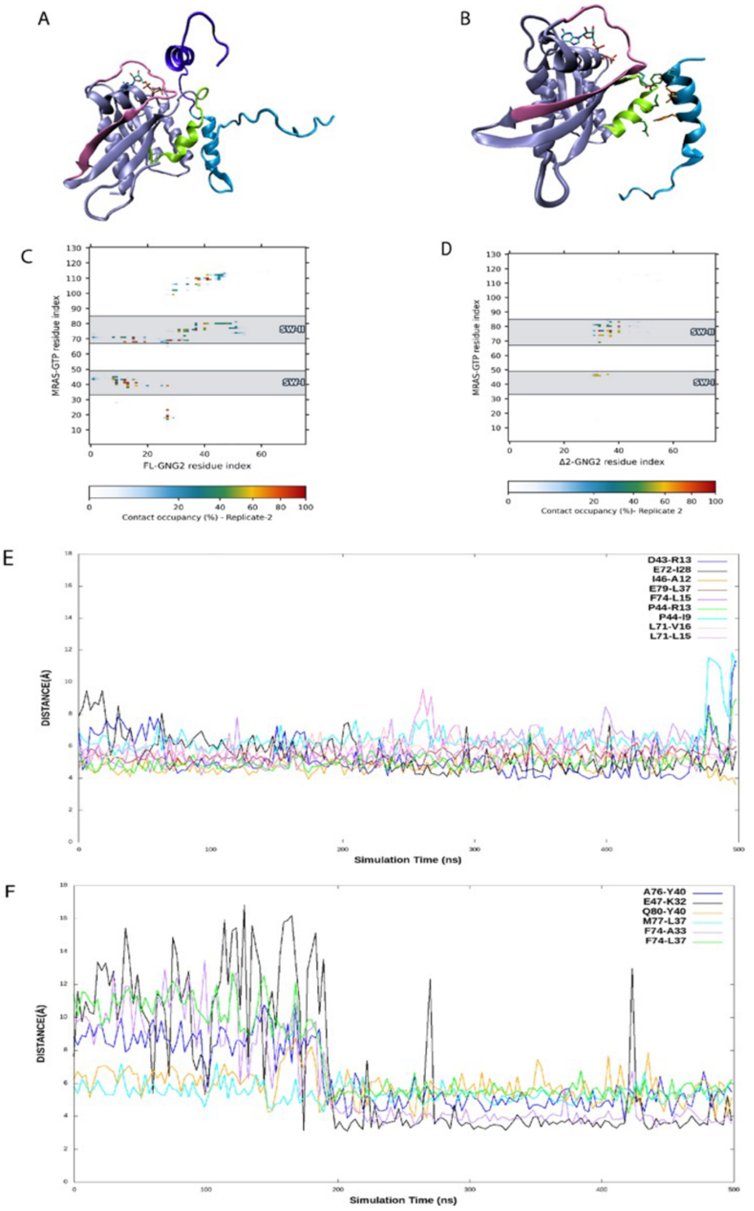


**FIGURE S5. Replicate of MD Simulation reveals GNG2 binding to the MRAS Switch I/II interface and competitively blocks SHOC2 binding**. **A**. Representative molecular dynamics (MD) simulation snapshot showing (GNG2-FL; purple-cyan) bound to the Switch I (mauve) and Switch II (lime) regions of MRAS (ice blue; PDB: 7TVF). GTP and $Mg^{2+}$ are displayed in licorice representation. **B**. Representative MD simulation snapshot of the GNG2-Δ2 peptide (cyan) bound to the Switch II regions of MRAS. **C, D.** Residue contact heat map of the MRAS FL-GNG2 and Δ2 complex. Contact maps were generated using a 5.0 Å heavy atom–heavy atom distance cutoff and demonstrate preferential peptide interactions with the Switch I and Switch II regions of MRAS. **E, F.** Distance plots demonstrating the stability of representative hydrophobic interactions, hydrogen bonds, and salt bridges between MRAS and (E) FL-GNG2 and (F) the GNG2-Δ2 peptide across the trajectory.

### Figure S6

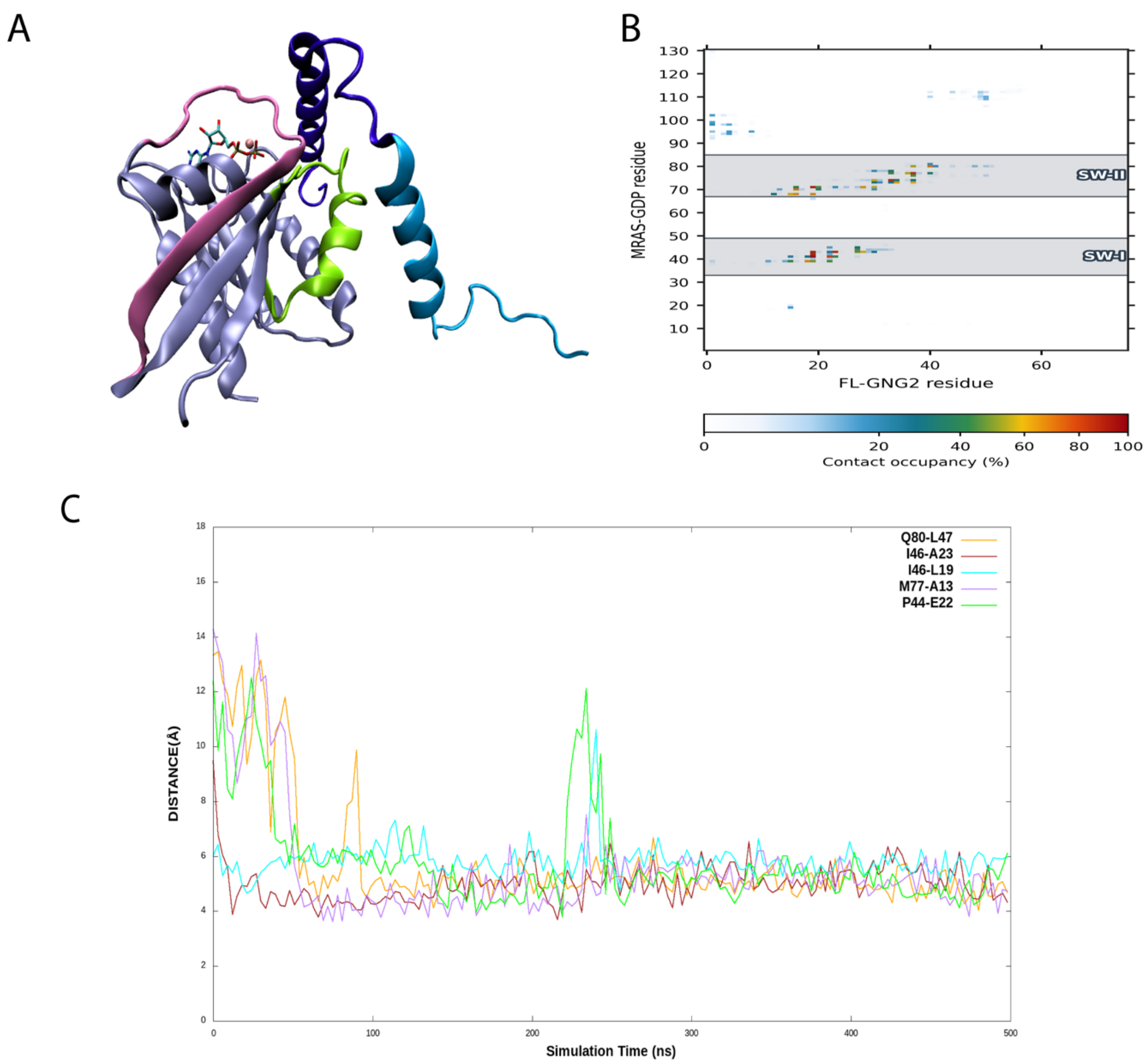


**FIGURE S6. GNG2 binding to the GDP- MRAS Switch I/II interface and competitively blocks SHOC2 binding surface**. **A**. Representative molecular dynamics (MD) simulation snapshot showing GNG2 full-length (GNG2-FL; purple-cyan) bound to the Switch I (mauve) and Switch II (lime) regions of MRAS (ice blue; PDB: (9C1A). GDP and $Mg^{2+}$ are displayed in licorice representation. **B.** Residue contact heat map of the MRAS FL-GNG2 Contact map was generated using a 5.0 Å heavy atom–heavy atom distance cutoff and demonstrate preferential peptide interactions with the Switch I and Switch II regions of MRAS. **C.** Distance plots demonstrating the stability of representative hydrophobic interactions, hydrogen bonds, and salt bridges between MRAS-GDP and FL-GNG2 peptide across the trajectory.

#### Figure S7

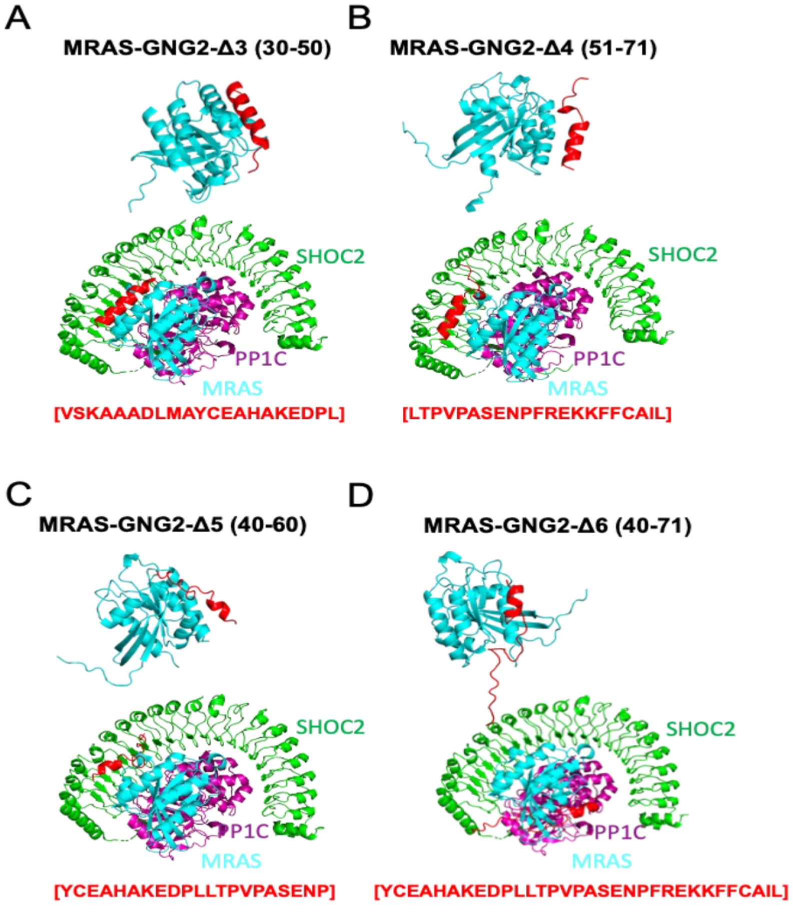


**FIGURE S7. AlphaFold2 models of GNG2-Δ2-derived variants bound to MRAS. A-D**. AlphaFold-predicted models of GNG2-Δ2-derived variants (Δ3–Δ6; red) bound to MRAS (cyan) and aligned with the SMP complex structure (SHOC2, green; PP1C, magenta; MRAS, cyan; PDB: 7TVF) to illustrate their predicted binding interfaces.

### Figure S8

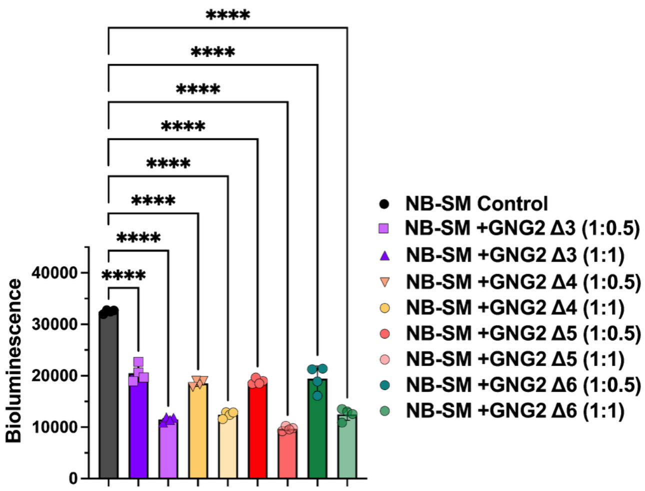


**FIGURE S8. GNG2-Δ2-derived variants inhibit the SHOC2–MRAS interaction in a dose-dependent manner.** Live-cell NanoBiT analysis showing SHOC2–MRAS interaction in HEK293 cells co-transfected with SHOC2$^{Lg}$-MRAS$^{Sm}$ plasmid along with small truncated Flag-GNG2 variants Δ3-Δ6 at a molar ratio of 1:0.5, and 1:1. Bioluminescence signals were measured 48 h post-transfection to assess the effect of GNG2 overexpression on the interaction of SHOC2–MRAS. Data are presented as mean ± S.D. (n = 4 biological replicates). Statistical significance was determined using one-way ANOVA comparing NanoBiT control and GNG2-expressing samples (***P ≤ 0.001, ****P ≤ 0.0001).

#### Figure S9

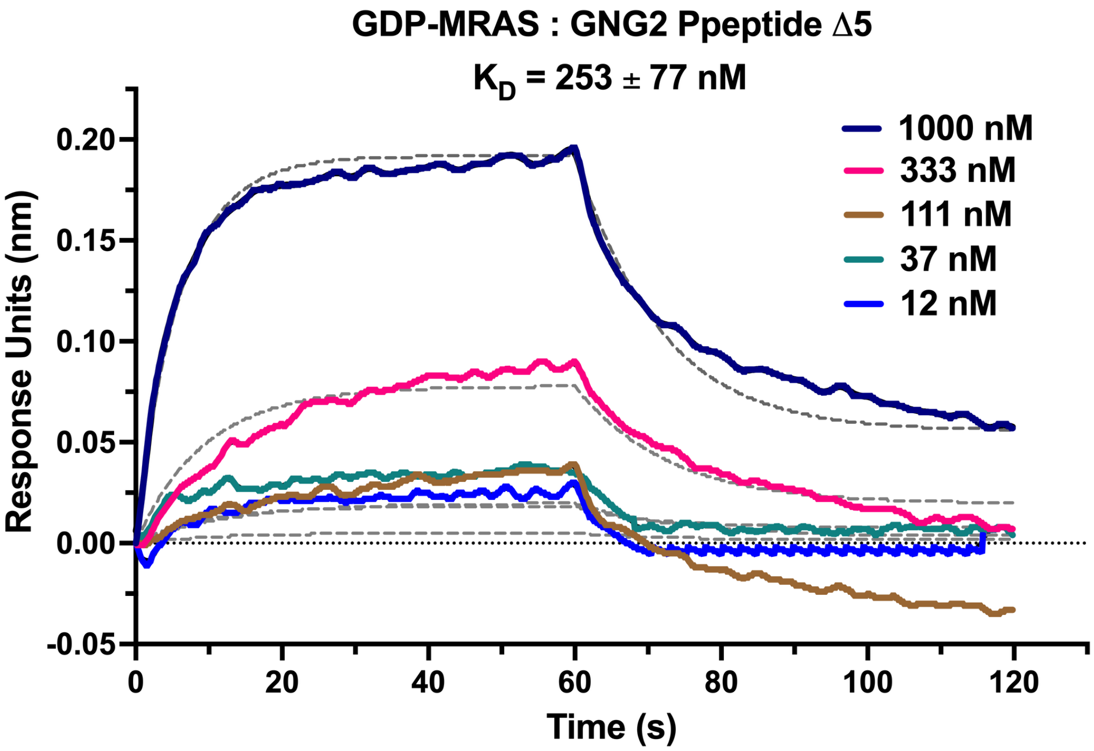


**FIGURE S9. TAT-conjugated GNG2 Δ5 peptide Interacts with GDP-MRAS.** Biolayer interferometry (BLI) analysis of the interaction between GDP-MRAS and GNG2 Δ5 peptide. GST-MRAS (GDP) was immobilized on GST biosensors, and increasing concentrations of GNG2 Δ5 peptide were used as analytes to determine binding kinetics. ($K_D$ of 253 ± 77 nM; $k_{on} = 1.95 \times 105\ M^{-1}s^{-1}$ and $k_{off} = 4.93 \times 10^{-2}\ s^{-1}$).
